# HIV-1 Vpr alters cellular transcription by targeting the PAF1 complex

**DOI:** 10.1101/2024.12.18.629146

**Authors:** Maximilien Biguet, Kelly Marno, Nidia Oliveira, Joseph M Gibbons, Corinna Pade, Christopher E Jones, Miguel R Branco, Aine McKnight

## Abstract

Innate immunity represents the first line of defence against viral infections, and successful pathogens like HIV-1 have evolved mechanisms to bypass these barriers. The PAF1 complex (PAF1c) has emerged as a key regulator of antiviral innate immune responses, inhibiting the replication of various viruses, including HIV-1. While its role in transcription regulation is well documented, little is known about how PAF1c inhibits HIV-1 and how HIV-1 circumvents this antiviral activity. Here we demonstrate that Vpr induces the rapid and transient downmodulation of PAF1c shortly after infection, a process that correlates with significant transcriptomic changes within the host cell. Transcriptomic profiling reveals that PAF1 knockdown mirrors many of the gene expression changes induced by HIV-1 infection, indicating that Vpr antagonism of PAF1c is a means by which HIV-1 may modulate host gene expression to favour replication. Further, we show that the loss of PAF1c leads to the suppression of interferon-stimulated genes (ISGs) and several known HIV-1 restriction factors. Overall, the evidence supports the emerging role of PAF1 as a regulator of a broad antiviral transcriptional program, but also suggests a specific anti-HIV activity that is countered by HIV-1 Vpr.

## INTRODUCTION

Innate immunity is the first line of defence that any invading virus must penetrate to establish infection and complete its replication cycle. Indeed, upon entering host cells, viruses trigger cascades of innate immune responses designed to restrict replication (Mesev et al., 2019). HIV-1 must therefore have evolved sophisticated strategies to breach this barrier, and understanding these mechanisms has become a critical focus of research (Laguette & Benkirane, 2015). Increasingly, attention is being directed towards uncovering how HIV-1 manipulates host innate immune pathways, as unravelling these interactions may provide valuable perspectives for advancing our understanding of HIV-1 pathogenesis and informing future treatment strategies (Colomer-Lluch et al., 2018; Ward et al., 2020).

The PAF1c is emerging as key regulator of antiviral innate immune responses. In 2011 we identified the PAF1c in a screen for early acting HIV-1 restriction factors (Liu et al., 2011). In the decade since, several groups have shown PAF1c to inhibit replication of other RNA viruses including Influenza (Marazzi et al, 2012), Dengue (Petit et al., 2021), and Zika (Shah et al., 2018). The PAF1c was initially discovered through its interaction with RNA polymerase II and has been shown to influence various processes associated with transcription (Van Oss et al., 2017; Francette et al., 2021). Transcriptional regulation by the PAF1c is increasingly understood to be context dependant as it has been found to either positively or negatively affect gene expression (He et al., 2011, Yu et al., 2015; Lu et al., 2016; Chen et al., 2015, Chen et al., 2017; Kenaston et al., 2022). Regulation of antiviral gene expression is thought to be the common mechanism by which it inhibits the replication of Influenza, Dengue, and Zika (Kenaston et al., 2022).

Although Liu et al. (2011) demonstrated that knockdown of PAF1c rescued HIV-1 infectivity, little is known of the mechanism by which PAF1c inhibits HIV-1 replication. PAF1c has been shown to affect the completion of HIV-1 reverse transcription (Liu et al., 2011) resulting in the production of fewer reverse transcripts, but also viral gene expression from the HIV-1 provirus LTR (Gao et al., 2020; Soliman et al., 2023). Given the known role of PAF1c in transcription regulation, here we hypothesise that PAF1c is also involved in regulating the early innate immune response to HIV-1 infection.

Following entry of HIV-1 into the cell, the viral RNA genome is reverse transcribed into double stranded proviral DNA for integration into host genomic DNA (Hu & Hughes, 2012). This reverse transcription step is influenced by other viral factors, such as the integrity of the viral capsid, which not only protects the viral genome but also regulates the timing and location of reverse transcription (Francis & Melikyan, 2018; Christensen et al., 2020). During these early stages of infection cellular proteins can sense the presence of these foreign nucleic acids, as well as certain viral proteins, and trigger signalling cascades resulting in the expression of antiviral genes such as interferon stimulated genes (ISGs) (Cheney et al., 2010; Yin et al., 2020).

HIV-1 has evolved to encode accessory proteins, which play a key role in evading this innate response (Laguette & Benkirane, 2015). Specifically, the accessory protein Vpr is a critical part of the virus’ arsenal to counter the early cellular innate immune response to infection (Desai et al., 2015; Bauby et al., 2021). Vpr is a small viral protein (14 kD) that is well conserved across retroviruses (Tristem et al., 1998) and plays a crucial role in facilitating efficient HIV replication, particularly in non-dividing cells like macrophages (Balliet et al., 1994; Eckstein et al., 2001; Mashiba et al., 2014). Vpr associates with the host DCAF1 E3 ubiquitin ligase complex and co-opts it to cause the ubiquitylation and subsequent proteasomal degradation of other host proteins (Zhao et al., 1994; Belzile et al., 2007; Lubow et al., 2020). Vpr is packaged in the virus particle and is rapidly transported to the nucleus upon infection, where it interacts with host proteins to promote viral replication (Paxton et al., 1993; Bachand et al., 1999; Desai et al., 2015). Although Vpr is one of the most extensively studied HIV-1 accessory proteins, pinning down its exact role in infection has remained challenging. One study suggested that Vpr may be causing changes in the levels of more than 2000 proteins (Greenwood et al., 2019). Given the small size of Vpr, it remains unclear how it would be able to interact directly with so many proteins. Instead, the evidence suggests many of these changes may not be a direct result of proteasomal degradation, Vpr may instead be targeting transcription factors to cause transcriptomic changes in the early stages of infection (Greenwood et al., 2019; Bauby et al., 2021; Reuschl et al., 2022). Indeed, Vpr appears to be solely responsible for most of the changes in gene expression observed within the first 12 hours of infection (Bauby et al., 2021), but also at later timepoints (Reuschl et al., 2022). Here we propose that PAF1c is one of the host transcriptional regulators targeted by Vpr, and that its downmodulation causes broad changes in host gene expression.

In this study, we investigate the interaction between HIV-1 and PAF1c. We demonstrate that Vpr induces the transient downmodulation of PAF1c shortly after infection, leading to broad transcriptomic changes. Indeed, we show that PAF1c knockdown by siRNA mirrors many of the transcriptomic changes observed during HIV-1 infection, suggesting that Vpr’s manipulation of PAF1c is a key driver of the host cell’s transcriptional response to the virus. Together, our findings reveal a novel mechanism by which HIV-1 exploits the host’s transcriptional machinery.

## RESULTS

### PAF1 Knockdown Rescues Infectivity of HIV-1 clinical isolates and in primary macrophages

To confirm the role of PAF1 in regulating HIV-1 replication, we first performed infectivity assays in CD4 positive HeLa cells treated with siRNA targeting PAF1 (siPAF1) or a control siRNA (siSCR). Four different clinical isolates of HIV-1, along with the 89.6 molecular clone, were used in these experiments. As shown in Figure 1A, knockdown of PAF1 significantly enhanced the infectivity of all viruses tested, as measured by focus-forming units per milliliter (FFU/mL) (p < 0.05). This observation was extended to primary human monocyte-derived macrophages (MDMs), where PAF1 knockdown also resulted in enhanced replication of the HIV-1 89.6 molecular clone (Figure 1B). PAF1 mRNA levels were found to be elevated in primary human monocytes, and to a lesser extent in Monocyte Derived Macrophages (MDMs), compared to HeLa-CD4 (Figure 1C).

**Figure 1:**
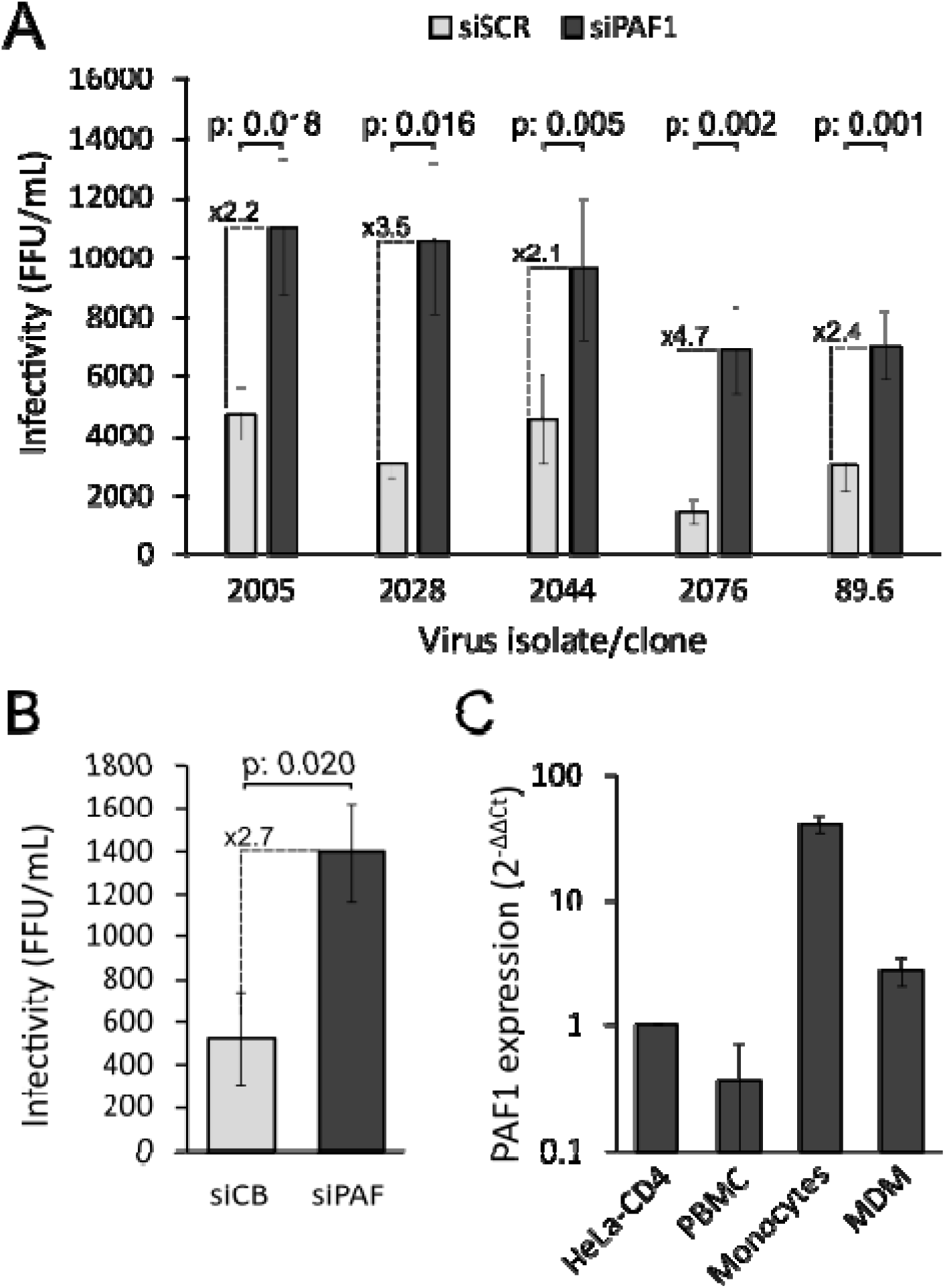
PAF1 knockdown rescues HIV-1 infectivity. **A.** Infectivity of four different clinical isolates of HIV-1 and the 89.6 molecular clone, measured in focus-forming units per milliliter (FFU/mL). The data represent paired infectivity measurements obtained from CD4/CXCR4 positive HeLa cells treated with a scrambled control siRNA (siSCR) or siRNA targeting PAF1 (siPAF1) across 5 replicates. p-values indicating significant differences in mean infectivity between control and PAF1 knockdown cells are shown (paired t-test, p < 0.05). Error bars = SEM. **B.** Infectivity of HIV-1 89.6 in primary human monocyte derived macrophages treated with a cyclophilin B control siRNA (siCB) or siRNA targeting PAF1 (siPAF) across 3 replicates, p-value shown above (paired t-test, p < 0.05). Error bars = SEM. **C.** β-actin normalized PAF1 mRNA expression levels in different primary human cells relative to HeLa-CD4 as determined by qPCR. Error bars: standard deviation. PBMC: Peripheral Blood Mononuclear Cells. MDM: Monocyte Derived Macrophages.

### HIV-1 Vpr Mediates Rapid Downmodulation of PAF1

We next sought to establish how HIV might escape from the antiviral action of PAF1. Imagestream analysis revealed a rapid reduction (-19%) in PAF1 protein levels within 30 minutes post-infection, reaching a maximum reduction at 90 minutes (-44%) post-infection before making a slight recovery (-35%) (Figure 2A, B). Western blotting confirmed that PAF1 protein levels declined as early as 30 minutes post-infection with the wild-type HIV-1 89.6 molecular clone but remained stable in cells infected with a ΔVpr virus (Figure 2C). Furthermore, infection of cells with Vpr-containing virus-like particles (VLPs) resulted in a similar downmodulation of PAF1 (Figure 2D). These observations show that HIV-1 Vpr is necessary and sufficient to induce the early loss of PAF1 protein levels.

**Figure 2.**
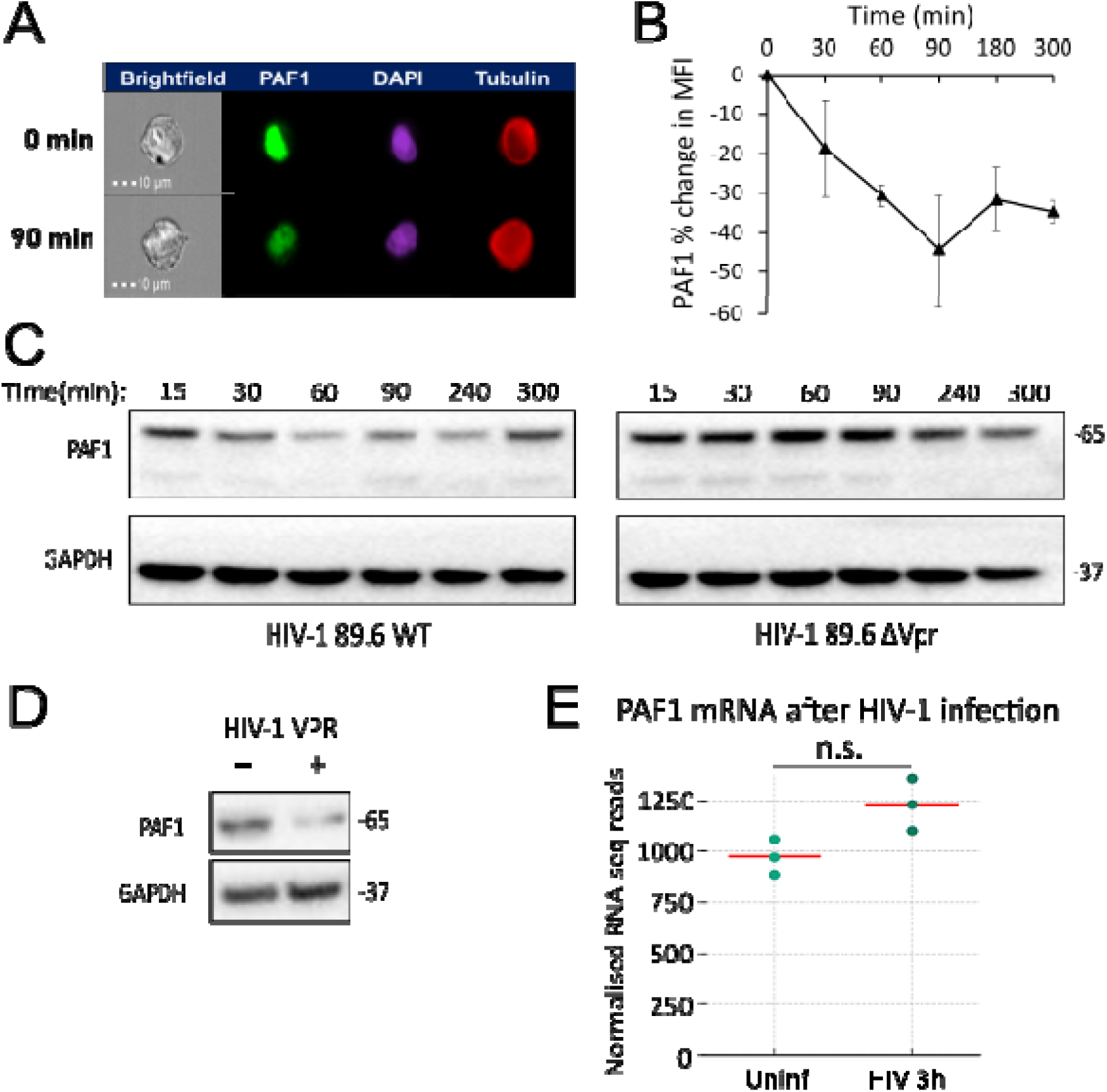
HIV-1 Vpr causes rapid loss of PAF1 protein levels early after infection. **A.** Representative pictures of imagestream analysis of PAF1 protein levels and subcellular location before infection and at 90 minutes post infection (MOI =1, infectivity determined by intracellular p24 staining). DAPI and tubulin used as nuclear and cytoplasmic markers respectively. **B.** Graph depicting the percent change in PAF1 MFI in HeLa-CD4 cells infected with the HIV-1 89.6 molecular clone (MOI =1). PAF1 levels were recorded by imagestream analysis over multiple timepoints covering the first four hours following infection. Data is derived from three replicates. Error bars: SEM. **C.** Representative image of a Western blot following PAF1 levels in HeLa-CD4 cells that were infected with either wild-type HIV-1 89.6 or ΔVpr (MOI = 1). PAF1 levels were recorded over multiple timepoints covering the first 24 hours following infection. The PAF1 signal is seen with a band at 65 kDa, and another at 90 kDa. GAPDH was used as a loading control. **D.** Representative image of a Western blot showing PAF1 levels in HeLa-CD4 cells treated with empty or Vpr-containing virus-like particles (VLPs). GAPDH was used as a loading control. **E.** Dot plot of DESeq2 normalised RNA-seq reads mapping to PAF1 transcripts in HeLa-CD4 cells infected with wild-type HIV-1 89.6 (3 hours post infection, MOI=1). The dots represent the read counts for each of three replicate experiments, the red horizontal line shows the mean of the normalized read counts for each condition. n.s.: not significantly differentially expressed.

Vpr is known to associate with the host DCAF-E3 ubiquitin ligase, co-opting it to mediate the ubiquitylation and subsequent proteasomal degradation of other host proteins (Zhao et al., 1994; Belzile et al., 2007; Greenwood et al., 2019). RNA-seq data shows that PAF1 mRNA levels remained unchanged at 3 hours post-infection, suggesting that the reduction in PAF1 protein is due to post-transcriptional mechanisms rather than decreased transcription (Figure 2E). This supports the idea that Vpr targets PAF1 for degradation without affecting its mRNA expression.

### PAF1 Interacts with REAF, Another Vpr Target

Previous studies have shown that HIV-1 Vpr also targets REAF, causing its down-modulation at similar timepoints after infection (Gibbons et al., 2020). Given this phenotypic similarity, and that both PAF1c and REAF are nuclear factors with roles in transcription (Winczura et al., 2021 - preprint), we hypothesized that these proteins may interact. In support of this, Western blot analysis of HeLa-CD4 cells treated with siRNAs targeting either PAF1 or REAF showed a concomitant reduction in the levels of both proteins (Figure 3A). Furthermore, qPCR data indicates that this reduction of REAF following PAF1 knockdown is not mediated by transcriptional regulation of REAF expression by the PAF1c (Figure 3B). Similarly, REAF siRNA knockdown does not cause a decrease in PAF1 mRNA levels (Figure 3B).

**Figure 3.**
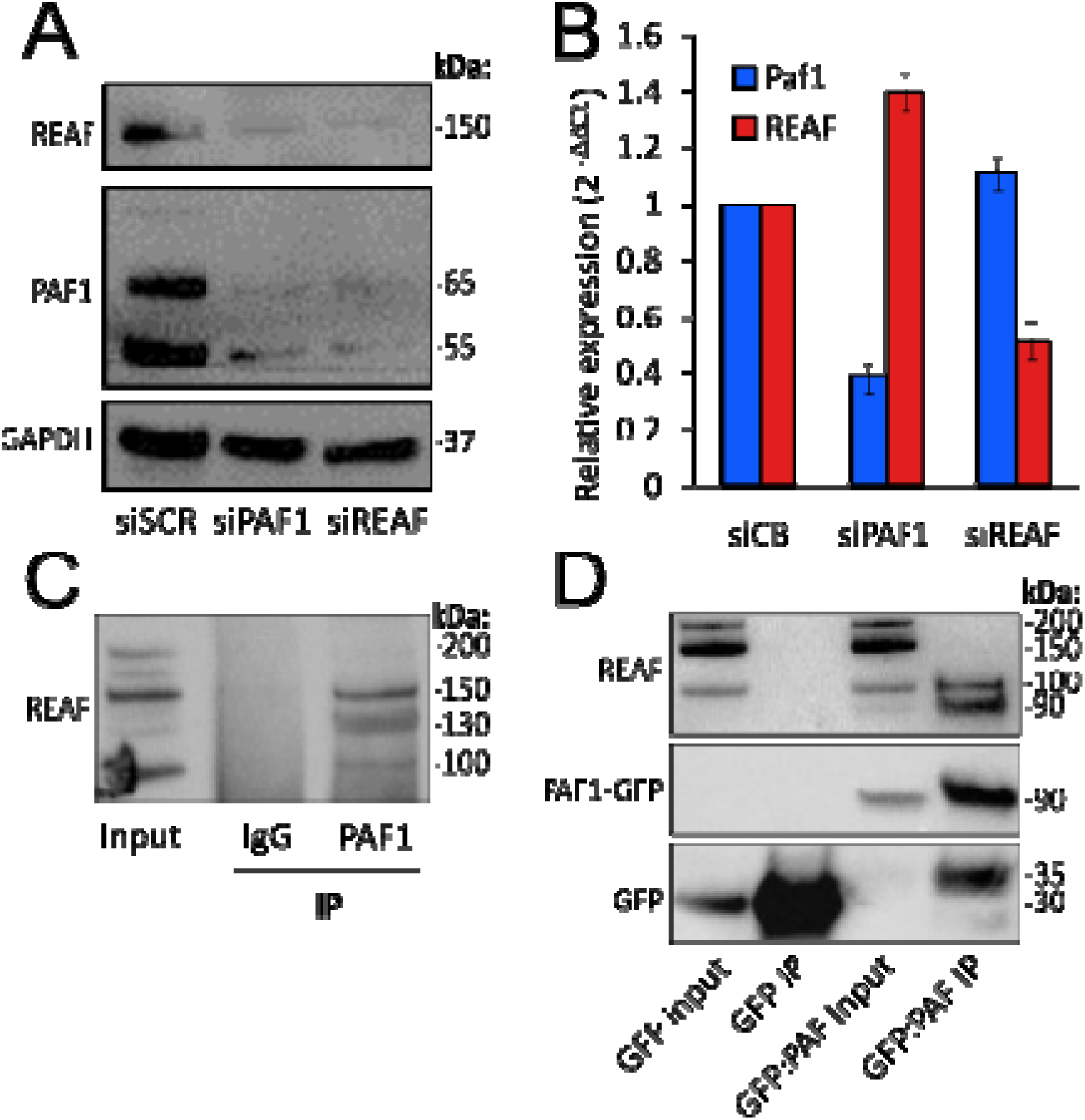
PAF1 interacts with REAF, another Vpr target. **A.** Western blot membrane showing REAF and PAF1 protein levels following treatment of HeLa cells-CD4 with a scrambled control siRNA (siSCR), PAF1 siRNA (siPAF1), or REAF siRNA (siREAF). GAPDH is included as a loading control. **B.** Representative qPCR data showing β-actin normalized PAF1 (blue) or REAF (red) mRNA expression levels after treatment with siRNA targeting PAF1 (siPAF1) or REAF (siREAF) relative to expression in cells treated with cyclophilin B control siRNA (siCB). Error bars: standard deviation. **C.** Endogenous PAF1 protein immunoprecipitation from a whole cell HeLa-CD4 lysate. Western blot analysis shows REAF protein levels in the input HeLa-CD4 cell lysate (input), following immunoprecipitation performed using an isotype control rabbit immunoglobulin (IgG IP), or following immunoprecipitation using a PAF1 antibody (PAF1 IP). Co-immunoprecipitated REAF was detected in the PAF1 immunoprecipitation only with band sizes of 100 and 150 kDa. **D.** HEK-293T cells were transfected with PAF1-GFP expression plasmid or GFP control vector. Western blotting was used to assess expression and immunoprecipitation with anti-GFP beads. Co-immunoprecipitated REAF was detected in the PAF1-GFP immunoprecipitation only, with a band at 100 kDa.

To confirm an interaction between PAF1 and REAF, we performed co-immunoprecipitation (Co-IP) experiments. PAF1 was immunoprecipitated from whole-cell lysates of HeLa-CD4 cells, and Western blot analysis revealed the presence of REAF in the PAF1 immunoprecipitate, REAF bands at 150 and 100 kDa map to known REAF isoforms (UniProt Consortium, 2023) (Figure 3C). Additionally, we transfected HEK-293T cells with a PAF1-GFP expression plasmid and used anti-GFP beads to pull down PAF1. Western blot analysis confirmed the co-immunoprecipitation of the 100 kDa REAF isoform with PAF1-GFP (Figure 3D). In contrast to the endogenous PAF1 immunoprecipitation, there was no REAF band at 150 kDa in the PAF1-GFP immunoprecipitation – suggesting an effect of the GFP:PAF1 fusion on this interaction with REAF. The 90 kDa band corresponds to PAF1-GFP.

### PAF1 Knockdown and HIV-1 Infection Induce Similar Transcriptomic Changes

Given the known role of PAF1c in regulation of host gene transcription (Van Oss et al., 2017, Francette et al., 2021), we investigated the transcriptomic changes associated with PAF1 knockdown and HIV-1 infection. We performed two RNA-seq analyses on HeLa-CD4 cells treated with siPAF1 or infected with HIV-1 89.6. The data shows that many of the genes regulated by HIV-1 infection at 3 hours (>30%) are also transcriptional targets of PAF1. Indeed, we identified 84 genes that were significantly regulated by both PAF1 knockdown and HIV-1 infection at 3 hours post-infection (Figure 4). 77 of the 84 common genes were concordantly regulated, their expression changing in the same direction (up or down) following either PAF1 siRNA knockdown or HIV infection. Gene ontology overrepresentation analysis (ORA) reveals there is an enrichment of transcriptional regulators in the list of 84 genes that overlap between our conditions (Supplementary file 1).

**Figure 4.**
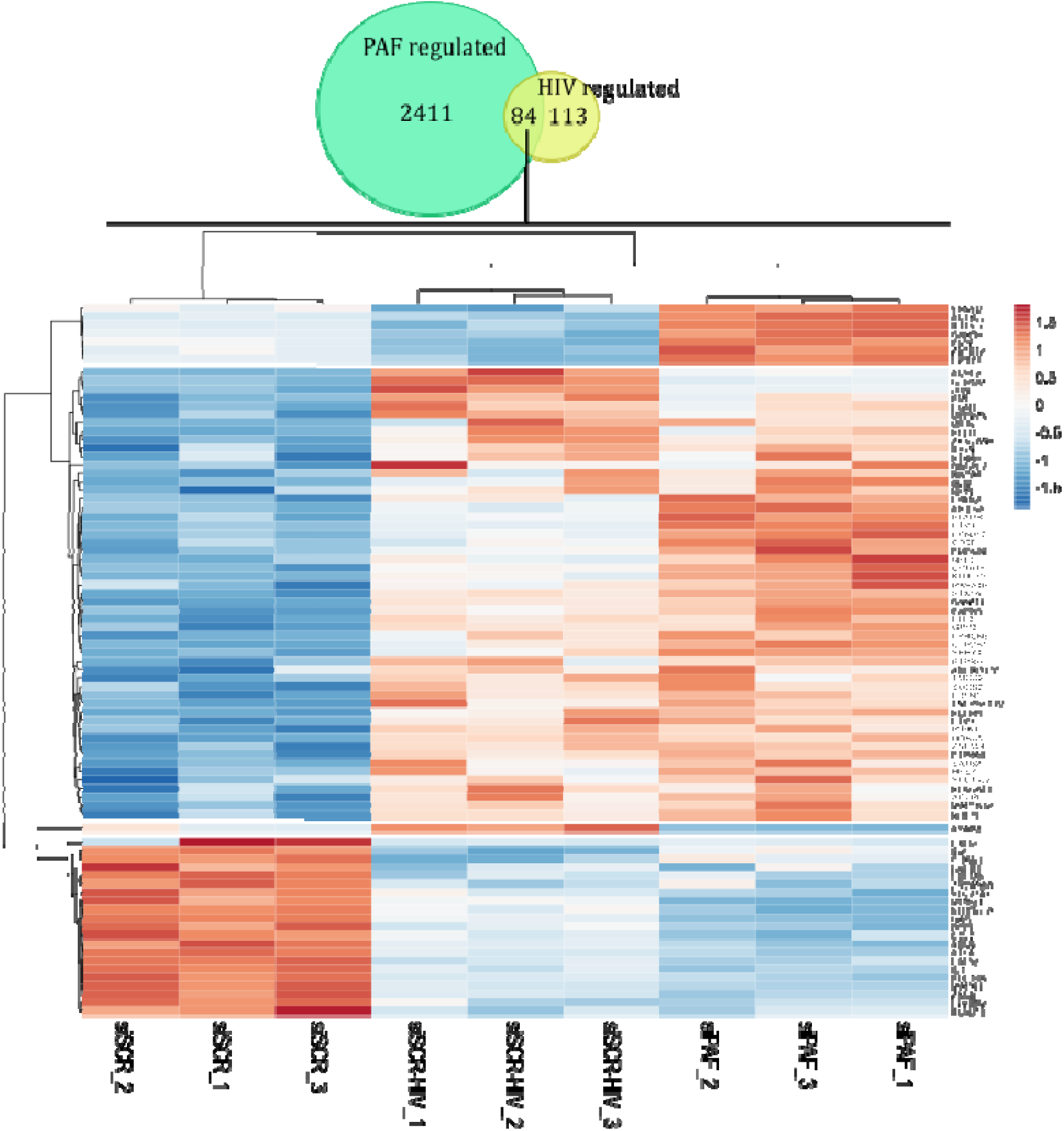
HIV-1 infection affects transcription of some PAF1 target genes. Heatmap showing expression of genes significantly regulated by both PAF1 knockdown and HIV-1 infection as determined by RNA-seq. 84 genes are significantly commonly regulated by PAF1 knockdown and HIV-1 infection (3 hours post infection, MOI = 1). A gene expression heatmap for these 84 genes was generated with variance stabilised expression data using ClustVis. Rows are divided into 4 clusters using correlation distance and average linkage shown by the adjacent dendrogram. Red/Blue scale indicates expression scores for each gene in each of the conditions specified at the bottom of the plot (siSCR = control siRNA treated, uninfected cells; siSCR-HIV = control siRNA treated, HIV-1 infected cells; siPAF1 = PAF1 siRNA treated, uninfected cells.

On a more global level of the transcriptome landscape, a custom gene set enrichment analysis (GSEA) also indicates that shifts, either up or down, in gene expression are essentially common between viral infection and siPAF1 knockdown (Figure 5A-D). An observed vs expected analysis was run on the overlap between PAF1 regulated genes and several sets of HIV-1 regulated genes (from either this study or Bauby et al., 2021). We find that each tested list of genes regulated by HIV-1 infection is significantly enriched for PAF1 target genes (p<0.05). However, only genes regulated at the earliest timepoints of HIV-1 infection (3h in this study, 4.5h in Bauby et al., 2021) are largely regulated in the same direction by PAF1 knockdown with 90% and 70% concordant regulation respectively. At later timepoints, despite PAF1 target genes being overrepresented in HIV-1 regulated gene sets, they are not necessarily regulated in the same direction. Overall, the data supports a model in which Vpr antagonism of PAF1 is a key mechanism by which HIV-1 alters host gene expression at an early stage of infection.

**Figure 5.**
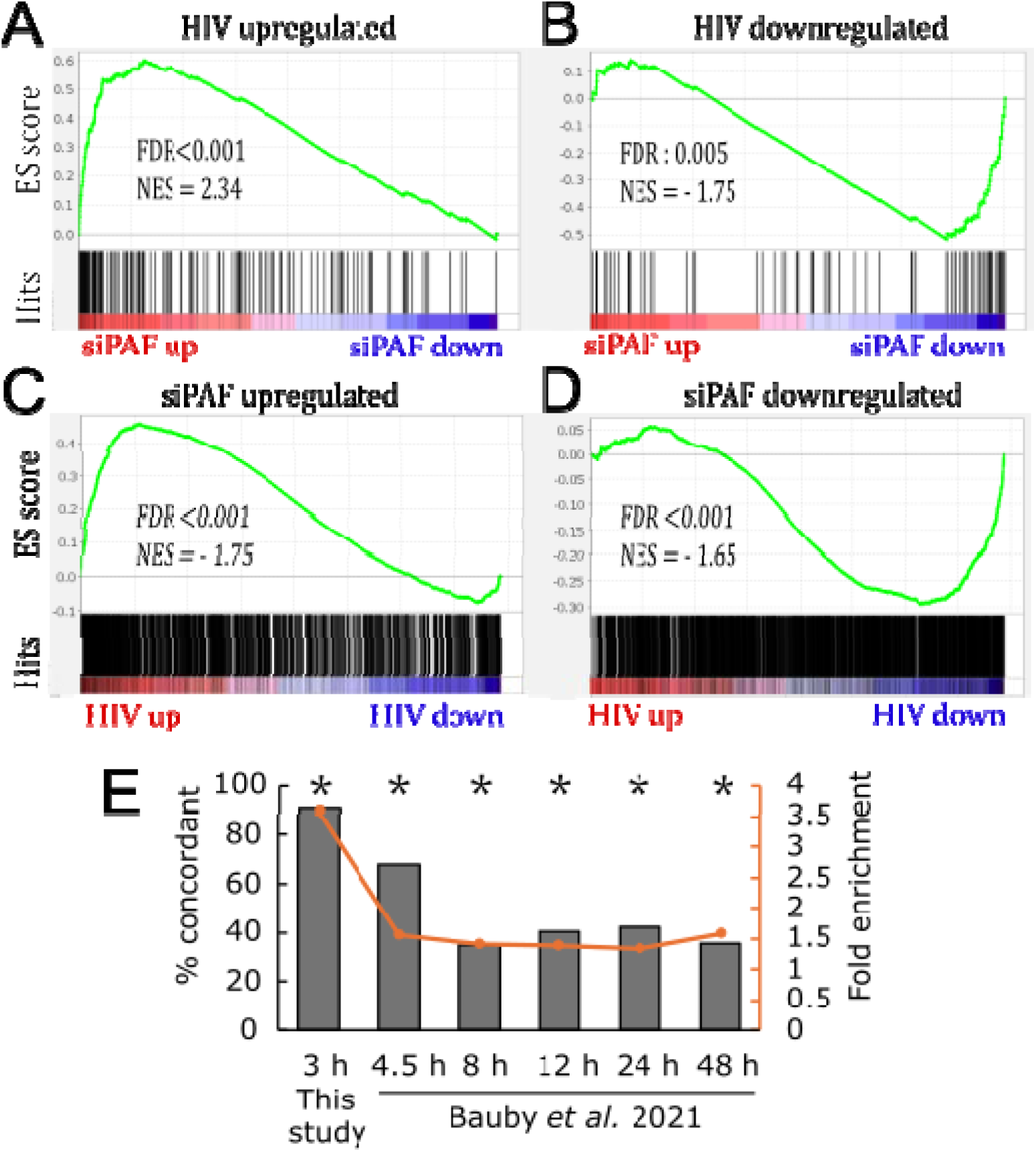
Genes broadly tend to be regulated similarly by PAF1 knockdown and HIV-1 infection at early timepoints. This figure presents several GSEA plots (**A-D**). For each gene-set named on top of the plots, the running enrichment score (ES) is shown as a green line. All genes are ranked by their expression following treatments detailed below (Red/Blue scale). Additionally, genes from the gene-set (referred to as ‘hits’) are marked as black bars positioned according to their respective places in the ranked list of all genes. The False Discovery Rate (FDR) and Normalised Enrichment Score (NES) are indicated for each analysis. Genes significantly upregulated by HIV at 3 hpi (**A**) or downregulated (**B**) were broadly dysregulated in the same direction by PAF1 siRNA treatment as evidenced by highly significant positive or negative enrichment respectively. Similarly, genes significantly upregulated by PAF1 siRNA knockdown (**C**) or downregulated by PAF1 knockdown (**D**) were broadly dysregulated in the same way by PAF1 siRNA treatment as evidenced by highly significant positive or negative enrichment respectively**. E.** Bar and line graph showing the results of an observed vs expected analysis on the overlap between PAF1 regulated genes and several sets of HIV-1 regulated genes on the x-axis. Left y-axis: percentage of genes in each overlap regulated in the same direction by PAF1 knockdown and HIV infection. Orange y-axis: fold enrichment (observed/expected number of genes that are significantly regulated by both PAF1 knockdown and each condition). Asterisks (**\***) represent FDR corrected p-values < 0.05 for a hypergeometric distribution test for statistical significance of the overlap size between each gene set and the PAF1 regulated gene set.

### PAF1 is a transcriptional regulator of immune genes

While many of the genes regulated by HIV-1 infection at 3 hours overlap with those affected by PAF1 siRNA knockdown, the overall scope of gene regulation is much broader in the knockdown condition after 72 hour treatment. PAF1 knockdown led to differential expression of nearly 2500 genes, compared with the 197 observed during early HIV-1 infection. We therefore interrogated this broader list of PAF1 transcriptional targets. We find that siRNA knockdown resulted in a widespread downregulation of interferon-stimulated genes (ISGs) and restriction factors, in line with its established role as a key transcriptional regulator of immune responses (Figure 6A, B). Gene Set Enrichment Analysis (GSEA) highlighted significant suppression of antiviral genes such as ISGs and known HIV-1 restriction factors including TRIM5, TRIM22, APOBEC3H, IFITM2, and SERINC5 following PAF1 siRNA knockdown (Figure 6C). Table 1 details all of the differentially expressed HIV-1 restriction factors following PAF1 siRNA knockdown and their fold changes. Flow cytometry analysis of cells expressing a GFP reporter under the control of an ISRE-mCMV promoter confirms that PAF1 is essential for the induction of ISG expression following poly(I:C) stimulation (Figure 6D, E, F).

**Figure 6.**
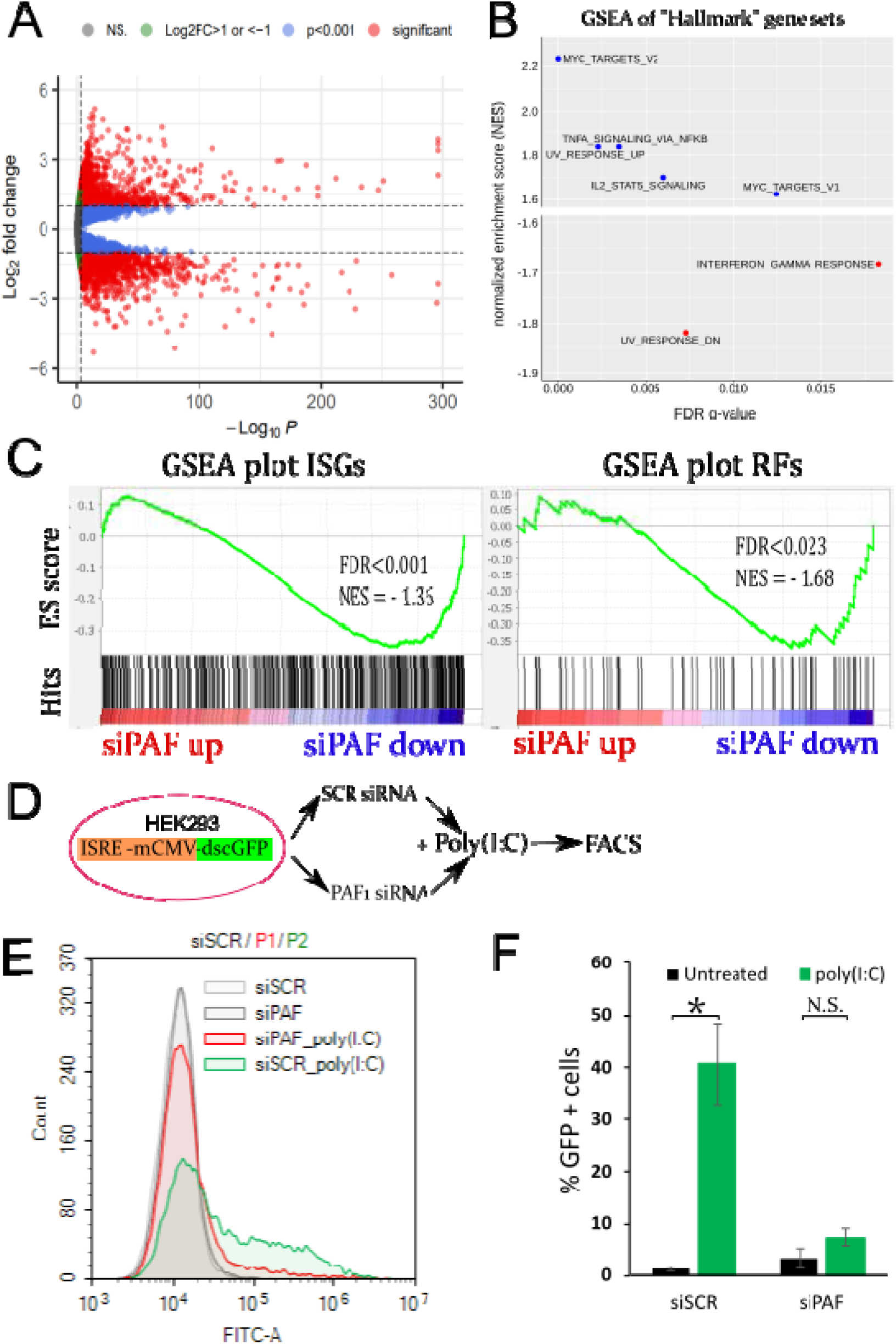
PAF1 is a transcriptional regulator of immune genes. **A.** Volcano plot of differentially expressed genes in HeLa-CD4 cells knocked down for PAF1. The Log2 fold change of mRNA levels shown on the Y axis is plotted against corrected p-value for genes. Genes that meet the selected significance and fold change cutoff (p<0.001 and Log2FC>1 or <-1) are coloured in red. **B.** Summary plots for GSEA analysis showing normalised enrichment score versus FDR q-value for with MSigDB Hallmark gene sets that are significantly upregulated (blue, NES>0) or downregulated (red, NES<0) following PAF1 siRNA treatment of cells. Significance cutoff was set at FDR < 0.05. **C.** GSEA plots for two curated gene sets (ISGs - genes with >10-fold induction on the interferome database of interferon stimulated genes; RFs – genes defined as restriction factors across peer-reviewed literature). The FDR and NES are indicated for each gene set on its respective plot. **D.** Diagram of the experimental setup and of the integrated reporter construct. The reporter cell line contains a dscGFP gene under control of an ISRE-mCMV promoter which can be activated with poly(I:C) treatment. dscGFP expression is measured by flow cytometry after stimulation of reporter cells transfected with either a control siRNA or a siRNA targeting PAF1. **E.** Histogram of GFP expression obtained by flow cytometry of the cell populations described in panel D and in the integrated legend. **F.** Percentage of GFP positive cells across the different tested conditions described in panel D. *: p-value < 0.05 for a student’s t-test.

**Table 1.** PAF1 transcriptionally regulates HIV-1 restriction factors. The table details the effect of PAF1 siRNA knockdown on expression of genes encoding for proteins described in the literature as HIV-1 restriction factors. Log2 FC/adj. p-value: log2 fold change of expression and FDR corrected p-value, respectively, for the given gene after differential expression analysis of PAF1 siRNA knockdown cells.

| Gene Name | log2 FC | adj. p-value | References |
| --- | --- | --- | --- |
| TRIM22 | -5.01 | 6.6E-31 | Tissot & Mechti., 1995; Kajaste-Rudnitski et al., 2011; Barr et al., 2008 |
| APOBEC3H | -2.98 | 4.4E-07 | Sheehy et al., 2002; Wang et al., 2011 |
| GBP2 | -2.91 | 7.8E-09 | Braun et al., 2019; Cui et al., 2021 |
| TRIM34 | -2.03 | 1.7E-25 | Ohainle et al., 2020 |
| LAPTM5 | -1.98 | 9.2E-04 | Zhao et al., 2021 |
| SHFL | -1.85 | 3.1E-43 | Wang et al., 2019 |
| TRIM5 | -1.32 | 1.3E-35 | Stremlau et al., 2004; Hatzioannou et al., 2004 |
| SERINC5 | -1.18 | 1.1E-10 | Usami et al., 2015 |
| IFITM2 | -1.06 | 2.4E-29 | Compton et al., 2014; Tartour et al., 2014 |
| ZC3H12A | 1.04 | 6.2E-19 | Liu et al., 2013 |
| TRIM28 | 1.05 | 9.2E-58 | Kamitani et al., 2008; Ait-Ammar et al., 2021 |
| ZC3HAV1 | 1.11 | 3.4E-64 | Ficarelli et al., 2021 |
| NOP2 | 1.14 | 1.1E-57 | Kong et al., 2020 |
| APOBEC3B | 1.69 | 2.5E-53 | Bishop et al., 2004 |

## DISCUSSION

This paper sought to delineate the interactions between HIV-1 Vpr and the PAF1c in modulating innate immunity. Our results show that HIV-1 Vpr antagonizes PAF1c, a transcription regulator of antiviral genes, by inducing its transient down modulation within an hour of infection making the cell more susceptible to superinfection during this phase. PAF1c siRNA knockdown after 48 hours results in loss of expression of antiviral genes including HIV-1 restriction factors and ISGs. We postulate that knockdown of PAF1 early in infection could have consequences in both early and late stages of replication.

PAF1c contains six proteins (Tomson & Arndt, 2013), and we previously observed that knockdown of any one component of the PAF1 complex enhanced infectivity of a reporter virus (Liu et al, 2011). We confirm this observation and extend it to several primary/clinical isolates of HIV-1. Additionally, we show that this effect also occurs in primary human macrophages, a key target cell of HIV-1 in which we find elevated PAF1 expression levels. PAF1c involvement in anti-viral cascades has been previously reported in defence against other viruses such as influenza, dengue and Zika (Marazzi et al., 2012; Shah et al., 2018; Petit et al., 2021). Strikingly, we see that PAF1 protein levels are transiently downmodulated by HIV Vpr in the first 30 to 240 mins of infection. It is known that Vpr associates with DCAF-E3 ligase and targets proteins for proteasomal degradation (Zhao et al., 1994; Belzile et al., 2007; Greenwood et al., 2019). Supporting the early timing of this event, Vpr is contained in the virus particle and rapidly localises to the nucleus where the majority of PAF1 is located (Desai et al. 2015).

We previously showed by immunoprecipitation that Vpr interacts with REAF and causes its degradation (Gibbons et al., 2020) and that REAF was a component of Lentiviral restriction 2 (LV2), an early restriction of HIV-1 infection (McKnight et al., 2001; Schmitz et al., 2004; Jackson-Jones et al., 2023). Here, we present evidence supporting the existence of a protein complex containing PAF1c components and REAF, leading us to postulate that PAF1c is another component of LV2 and that Vpr antagonises this whole protein complex.

By targeting transcription factor PAF1, our observations support the possibility that Vpr activity could additionally have later effects on subsequent events in HIV replication by modulating host gene expression. We show that HIV-1 infection causes transcriptomic changes at 3 hours post infection, and that many of the differentially expressed genes are similarly affected by PAF1 siRNA knockdown and HIV-1 infection. A recent study by Bauby et al. (2021) demonstrated that targeting by Vpr of a yet unknown host factor is the cause of the vast majority of transcriptomic changes observed in the first 12 hours of infection. Our results suggest that antagonism of PAF1c by HIV-1 Vpr is at least partially responsible for this early manipulation of the cellular transcriptome by HIV-1 infection. This is supported by previous study on Vpr activity (Bauby et al., 2021). Indeed, despite significant experimental differences, we find that PAF1 target genes are overrepresented in the list of genes Bauby et al. find to be regulated by HIV-1 infection at various timepoints. Furthermore, at the earliest timepoint tested (4.5 hpi) genes were broadly regulated in the same direction by PAF1 knockdown or HIV-1 infection. Overall, strong parallels can be drawn between HIV-1 infection and loss of PAF1. This makes sense from an evolutionary perspective where targeting key transcriptional regulators such as PAF1c may be a means by which HIV-1 and other viruses can more efficiently modify the cellular environment avoiding the task of counteracting thousands of individual host factors by directly targeting them. This is supported by the apparent functional convergent evolution of viral accessory proteins across several virus families (Flaviviridae, Orthomyxoviridae) to target PAF1c and its transcriptional activity (Marazzi et al., 2012; Shah et al., 2018; Petit et al., 2021). For example, the influenza protein NS1 acts as a histone mimic and interacts with PAF1c to prevent it from inducing ISG expression (Marazzi et al., 2012).

The specificity afforded by targeting PAF1c is surprising. Our RNA-seq analysis reveals broad transcriptional changes upon PAF1c knock-down with siRNA in HeLa-CD4 cells after 48 hours. On the other hand, infection of HeLa-CD4 cells with HIV-1 resulted in only 197 genes differentially expressed by 3 hours suggesting that PAF1c targeting by Vpr is more specific than knockdown by siRNA. It will be interesting to test a potential mechanism that only PAF1c controlled genes that are also bound by REAF are targeted. We show that PAF1c knock-down by siRNA causes a broad downregulation of antiviral factors, including ISGs and several known HIV-1 restriction factors. We also show that PAF1c is necessary for the induction of ISG expression following poly(I:C) stimulation supporting the key role of PAF1c in the activation of anti-viral responses.

It is known the PAF1c downmodulation can result in either up or down modulation of gene transcription (Kenaston et al., 2022). Potentially PAF1c downmodulation could promote infection by, in addition to depleting anti-viral factors, inducing expression of genes that help viral replication either directly or those that negatively regulate anti-viral genes. Indeed, among the genes found to be positively regulated by PAF1 siRNA knockdown and HIV-1 infection, several have already been shown to be involved in assisting HIV-1 replication. A genome-wide screen for positive host factors required for the early steps of infection found that knockdown of AT-rich interactive domain 5A (ARID5A) inhibited replication of a VSV-G pseudotyped HIV-1 in HEK-293T cells (Konig et al 2008). Furthermore, three more factors: pellino E3 ubiquitin protein ligase 1 (PELI1), lysophosphatidic acid receptor 2 (LPAR2), and oxysterol binding protein like 7 (OSBPL7) that enhance HIV-1 replication (Zhou et al., 2008) are shown here to be significantly induced upon both PAF1 siRNA knockdown and HIV-1 infection at 3 hours. Finally, both PAF1 siRNA knockdown and HIV-1 infection (at 3 hours) upregulate JUN, a component of the AP-1 transcription factor reported to be upregulated and activated during HIV-1 infection (Duverger et al., 2013). AP-1 is involved in enhancing transcription of viral genes from the LTR (Colin & Van Lint, 2009). Thus, antagonism of PAF1c may be one mechanism by which HIV-1 can manipulate JUN expression.

We previously reported that PAF1c and REAF inhibit the reverse transcription step of HIV-1 replication (Liu et al., 2011; Gibbons et al., 2020). While we cannot rule out that PAF1c and REAF may directly affect the process of reverse transcription, our findings support a role for PAF1c in regulating the expression of genes that could affect various stages of the viral life cycle including reverse transcription and events surrounding it such as capsid integrity. We find that HIV-1 Vpr antagonism of PAF1c facilitates viral replication by altering the cellular transcriptome. Future studies will be essential to unravel the full scope of Vpr’s influence on host transcription.

## METHODS

### Cell culture

All cell lines were cultured at 37°C in a 5% CO_2_ atmosphere, regularly monitored and passaged when they reached 80-90% confluency. CD4/CXCR4 positive HeLa cells (HeLa-CD4, NIBSC) were cultured in High Glucose Dulbecco’s Modified Eagle Medium (DMEM, Gibco CN: 10566016), enriched with 5% heat inactivated foetal calf serum (FCS, Sigma Aldrich CN F7524) and antibiotics (100 units/mL penicillin and 100 μg/mL streptomycin), G418 antibiotic was used for purifying selection of CD4 expressing cells. HEK-293T and ISRE-293 reporter cells were grown with High Glucose DMEM supplemented with 10% FCS. The ISRE-293 reporter cell line was kindly gifted to us by J. Rehwinkel (see Hertzog et al 2018). Adherent cell lines were sub-cultured by washing with Dulbecco’s phosphate buffered saline (PBS, Sigma CN: D857) prior to a 5-minute incubation with a 0.05% Trypsin-EDTA solution (Gibco CN: 15400054). C8166 cells (NIBSC) were cultured in RPMI-1640 medium (Gibco CN: 11875093) supplemented with 10% FCS, 2 mM L-glutamine (Gibco CN: A2916801), and antibiotics.

### Virus and Virus-Like Particle (VLP) Production

HEK-293T cells were transfected to produce HIV-1 89.6 molecular clone virus (NIBSC). Cells were seeded in 6 well plates with antibiotic-free DMEM containing 10% FCS and cultured to 80-90% confluency. For each well, 3 µg of HIV-1 89.6 plasmid DNA was transfected using lipofectamine 3000 (Invitrogen) as per manufacturer’s instructions. After overnight incubation, the medium was replaced with DMEM supplemented with 10% FCS, and the cells were incubated for 48 hours. HEK-293T cells were then scraped and co-cultured with a confluent C8166 cell culture in complete RPMI-1640. The mixture was centrifuged at 500 g and incubated for 1-2 hours at 37°C to facilitate infection. After resuspension in 25 mL of complete RPMI, the cells were cultured at 37°C with 5% CO_2_. After 48 hours, the supernatant was collected, centrifuged at 1000 g to remove debris, and stored at -80°C.

LPEI transfection was used for VLP production with a 1:1:1 mass ratio of plasmids for VSV-G expression (Stratagene CN: 217567), packaging vector, and HIV-1 VPR expression. The packaging vector and VPR expression vectors were gifts from N. Landau (Connor et al., 1995). Control VLPs were generated by replacing the VPR plasmid with pcDNA plasmid. A total of 24 µg of plasmid DNA was transfected in an 80% confluent 10 cm cell culture dish, and the media was refreshed after 24 hours. VLP-containing supernatant was harvested at 48 and 72 hours post-transfection, centrifuged at 1000 g, and stored at -80°C.

### Infectivity Assay for Replication-Competent Virus

Infectivity was measured using a focus-forming unit assay (McKnight et al., 1994). HeLa-CD4 cells (1.4 x 10^4) were seeded in 48-well plates 24 hours before infection. Serial dilutions of virus stock were added in 200 µL volumes per well. After 48 hours of incubation, with a media change at 24 hours, cells were washed with PBS and fixed with ice-cold methanol/acetone (1:1) for 10 minutes. Fixed cells were stained with mouse anti-p24 antibody (MRC AIDS Reagent Program UK, EVA 365 and 366) (1:50) for 1 hour at 37°C, followed by β-galactosidase-conjugated goat anti-mouse antibody (Southern Biotech, 1010-06) (1:500) for another hour. Cells were washed, and β-galactosidase substrate was added before overnight incubation at 37°C. Foci of infection were counted to determine virus stock titre.

### HIV-1 p24 ELISA for VLP Titration

HIV-1 p24 levels in VLP stocks were quantified using a sandwich ELISA. Samples were prepared by mixing 87 µL of virus-containing supernatant with 3 µL of 30% Empigen BB detergent (Sigma-Aldrich, 66455-29-6), followed by heat inactivation at 56°C. Samples were diluted with TBS containing 0.1% Empigen BB. Standards were prepared by diluting purified p24 antigen (Abcam, ab127888) in TBS with Empigen BB. High-affinity ELISA plates (PerkinElmer, 6005500) were coated overnight with sheep anti-HIV-1 p24 antibody (Aalto, D7320) in sodium bicarbonate buffer at 4°C. Plates were blocked with 5% BSA in TBS, and samples or standards were incubated for 3 hours at room temperature. After washing, alkaline phosphatase (AP)-conjugated anti-HIV-1 p24 antibody (Aalto, BC 1071-AP) was added (0.5 µg/mL) and incubated for 1 hour. Plates were washed and incubated with AP substrate and Sapphire II enhancer (ThermoFisher, T2210), and chemiluminescence was measured using a plate reader. p24 concentrations were calculated by comparison to the standard curve.

### PAF1 Knockdown via siRNA

HeLa-CD4 cells were transfected with PAF1, REAF, or control siRNA (Qiagen) using HiPerfect reagent (Qiagen) in 6-well plates. A siRNA/lipid complex was prepared by mixing 190 µL Opti-MEM, 8 µL HiPerfect, and 3 µL siRNA (20 µM stock) and incubated for 30 minutes. HeLa-CD4 cells (63,000 per well) were plated in DMEM with 10% FCS (no antibiotics) and the complex was added. After overnight incubation at 37°C, media was replaced with fresh DMEM, and cells were incubated for 48 hours. For ISRE-293 cells, siRNA was diluted in Opti-MEM (100 nM final concentration) and mixed instead with DharmaFECT1 (Dharmacon) before adding to cells (1.5 x 10^5 per well). Cells were incubated for 48 hours at 37°C before further use.

### Flow cytometry & Imaging Flow Cytometry

GFP expression in ISRE-293 cells was measured using a NovoCyte flow cytometer (Agilent). Cells were trypsinized, pelleted at 1000 g, washed with PBS, and fixed with Fix and Perm solution A for 15 minutes. After two PBS washes, cells were resuspended in PBS with 5 mM EDTA and 5% FCS, then loaded onto a 96-well plate for analysis.

For imaging flow cytometry, cells were collected by trypsinization, pelleted at 1000 g, and washed with PBS. They were fixed with Fix and Perm solution A (Nordic MUbio) for 30 minutes, then permeabilized with 0.2% Triton X-100 in PBS for 20 minutes. Cells were incubated at 4°C for 24 hours with 2 µg/mL primary antibody in staining buffer (PBS, 0.1% Triton X-100, 0.5% FBS). After washing, cells were stained with 1 µg/mL fluorophore-conjugated secondary antibody for 1 hour in the dark and then washed again. Nuclei were stained with 1 µg/mL DAPI for 2 hours before a final wash. Samples were analyzed using an ImageStreamX Mark II Flow Cytometer (CYTEK), capturing images for 5000 cells per sample. Data analysis was performed with IDEAS software.

### Western Blotting

Western blotting was used to analyze protein expression. Cells were harvested by trypsinization, pelleted at 500 g, washed with PBS, and lysed with RIPA buffer (50 mM Tris-HCl, pH 7.4, 150 mM NaCl, 1% NP-40, 0.5% sodium deoxycholate, 0.1% SDS) containing protease and phosphatase inhibitors (Thermo Scientific). Lysates were incubated on ice for 15 minutes and then centrifuged at 12,000 g to remove debris. Protein concentration was measured using the Pierce BCA kit (Thermo Scientific). Equal amounts of protein (10 µg per lane) were denatured at 95°C for 5 minutes and separated by SDS-PAGE using 4-12% Bis-Tris gels (Invitrogen) in MES-SDS buffer. Proteins were transferred onto a PVDF membrane using wet transfer at 50 V for 2 hours. The membrane was blocked in 5% non-fat milk in TBST for 1 hour and incubated overnight at 4°C with primary antibodies. After washing, the membrane was incubated with HRP-conjugated secondary antibodies for 1 hour. The signal was detected using the SuperSignal West Pico Plus ECL system and imaged with a ChemiDoc MP system (BioRad).

### Immunoprecipitation

For immunoprecipitation, 2 x 10^6^ cells were harvested, washed with PBS, and pelleted at 500 g. Cells were lysed in immunoprecipitation buffer (50 mM Tris pH 7.5, 5 mM MgCl2, 150 mM NaCl, 10% Glycerol, 1% NP-40) with protease and phosphatase inhibitors for 15 minutes on ice. Lysates were cleared by centrifugation at 12,000 g, and protein concentration was determined using the BCA assay. For GFP-tagged proteins, GFP-trap magnetic beads (ChromoTek) were used according to manufacturer instructions. For endogenous proteins, 25 µL of protein A magnetic beads (Invitrogen) were incubated with 5 µg of antibody in 200 µL PBS with 0.05% Tween-20 for 1 hour at 4°C. After washing, the bead-antibody complexes were incubated with 500 µL of lysate containing 1 µg protein for 3 hours. Beads were washed three times with PBST, resuspended in 2X Bolt LDS buffer, and boiled at 95°C for 5 minutes before Western blotting.

### Antibodies

For imaging flow cytometry, the following antibodies were employed: rabbit polyclonal anti-PAF1 (Proteintech, 15441-1-AP) and rat monoclonal anti-tubulin-α (Abcam, ab6160). Detection was achieved using Alexa Fluor 488-conjugated donkey anti-rabbit IgG (Invitrogen, A21206) and Alexa Fluor 647-conjugated goat anti-rat IgG (Invitrogen, A21247). For Western blot analysis, the primary antibodies used were: rabbit polyclonal anti-PAF1 (Proteintech, 15441-1-AP), rabbit polyclonal anti-REAF/RPRD2 (Eurogentec, Marno et al., 2014), rabbit monoclonal anti-IFITM3 (Insight Biotechnology), rabbit polyclonal anti-GAPDH (Abcam, ab9485), and rabbit polyclonal anti-GFP (Proteintech, 50430-2-AP). Secondary detection was carried out using horseradish peroxidase-conjugated goat anti-rabbit IgG (Invitrogen, A16096). For intracellular staining of HIV-1, a monoclonal antibody against p24 (MRC AIDS Reagent Program UK, EVA 365 and 366) was used, followed by detection with a goat anti-mouse IgG β-galactosidase-conjugated antibody (Southern Biotech, 1010-06). For ELISA, we utilized an affinity-purified anti-HIV-1 p24 antibody (Aalto, D7320) along with alkaline phosphatase-conjugated anti-HIV-1 p24 (Aalto, BC 1071-AP).

### RNA-sequencing and data analysis

Cells were collected by trypsinisation and pelleted in 1.5 mL microcentrifuge tubes, they were washed once with 1 mL PBS then pelleted and stored at -80°C. Samples were later processed in a single batch using an RNeasy mini RNA purification kit (Qiagen CN: 74104). RNA was resuspended in DEPC treated water; genomic DNA contamination was removed using the Turbo DNA-free removal kit (Invitrogen CN: AM1907). RNA quality and quantity assessed by nanodrop spectrophotometry to ensure it matched sample submission requirements for RNA-sequencing services by Novogene (Illumina library preparation with polyA enrichment, paired end 150 bp RNA-sequencing on NovaSeq instrument). Each library was sequenced to a depth of 20 million reads.

All analysis of next generation sequencing data utilised Queen Mary University’s Apocrita high performance computing facility, supported by QMUL Research-IT (Thomas et al., 2017). First, adapters (auto detected) and low-quality bases (phred score < 20) were trimmed and reads shorter than 25 bp were discarded using Cutadapt (Martin, 2011). Gene expression levels were quantified using RSEM (Li and Dewey, 2011) and its integrated Bowtie2 alignment (Langmead & Salzberg, 2012). Reference genome assembly files for human build GRCh38 version 45 were downloaded from the GENCODE website (https://www.gencodegenes.org/human/). DESeq2 (Love et al., 2014) was used for differential expression analysis with normal parameters. Genes with an adjusted p-value < 0.001 and a log2 fold change >1 or <-1 were considered to be differentially expressed.

Gene set enrichment analysis was performed using the Broad Institute’s GSEA tool (Subramanian et al., 2005; Mootha et al., 2003). DESeq2 normalised reads were used as input, and GSEA analysis was run using various gene-set databases (MSigDB – Liberzon et al., 2011; REACTOME – Milacic et al., 2024; Fabregat et al., 2017) and custom gene-sets as described in the relevant sections. The GSEA tool was then run using the gene-set permutation mode with 1000 permutations.

## Notes

### Competing Interest Statement

The authors have declared no competing interest.

### Summary of Updates

Corrected author list due to missing author

